# Exploring the deletion landscape of *S. aureus* Cas9 with SABER

**DOI:** 10.1101/2025.08.12.669962

**Authors:** Andrew J. Plebanek, Luke M. Oltrogge, Cynthia I. Terrace, Maria Lukarska, Maggie Khoury, David F. Savage

**Affiliations:** Department of Molecular and Cell Biology, University of California, Berkeley, Berkeley, CA 94720, USA; Innovative Genomics Institute, University of California, Berkeley, Berkeley, CA 94720, USA; Howard Hughes Medical Institute, University of California, Berkeley, Berkeley, CA, 94720, USA; School of Pharmacy, University of California, San Francisco, San Francisco, CA 94143

## Abstract

Profiling tolerated amino acid deletions in proteins can elucidate structure-function relationships, reconstruct intermediate stages in protein evolution, and be used to engineer minimized versions of proteins with size-sensitive biotechnology applications. Despite advances in deletion library construction techniques over the past several decades, there are presently few methods available that are simultaneously efficient, precise, and easy to implement. Here we present SABER, a novel approach which utilizes SpRYCas9, a near-PAMless engineered SpCas9 variant, as a molecular biological tool for building deletion libraries with unprecedented speed and ease. We applied this technique to the small and structurally divergent Cas9 from Staphylococcus *aureus* (SaCas9) and mapped the set of deletions tolerated for DNA binding activity. We proceeded to use this information to design a set of minimal SaCas9-based effectors capable of CRISPRi transcriptional repression in bacterial cells. Our findings provide new insights into the function of certain structural elements in SaCas9, and we anticipate that our dSaCas9 deletion map may prove useful in further efforts to develop minimal Cas9-based effectors and gene editors.

## Introduction

Proteins evolve by accumulating mutations, which may take the form of either changes in the identity of a single amino acid residue at a given position (substitutions), or topological changes typically involving the insertion or deletion of one or more amino acids (indels). Within this latter category, deletions are broadly more common than insertions in organismal genomes from across the tree of life (Gregory 2004; Taylor et al. 2004; Savino et al. 2022), and in protein-coding genes may range in size from omissions of single codons to the loss of entire domains (Weiner et al. 2006). In most cases, deletions are associated with a significant fitness cost, as they may not only remove key residues or disrupt the open reading frame, but can also result in broader structural changes that may significantly impact protein folding and function (Savino et al. 2022). However, in certain contexts deletions may be neutral or even beneficial with respect to protein fitness, potentially enhancing enzymatic activity or contributing to the evolution of novel activities (Patzoldt et al. 2006; Yi et al. 2012; Hwang et al. 2014; Yang et al. 2019).

Reducing the size of a protein by deleting amino acids may also be desirable in the case of certain proteins with size-sensitive biotechnology applications. Perhaps the foremost example of this may be seen in Cas9 and other RNA-guided DNA-binding proteins, several of which have been adopted for *in vivo* genome editing applications using viral delivery to mammalian tissues, but which are often too large for their genes to be effectively packaged into commonly-used viral vectors (Doudna et al. 2020). The large size of these effectors becomes especially problematic in cases where additional protein domains are fused to the effector module, as in base- and prime-editing Cas-fusion constructs (Anzalone et al. 2020). In response to this challenge, researchers have drawn upon metagenomics datasets to identify novel effectors small enough to be effectively incorporated into viral vector genomes (Burstein et al. 2017). However, these effectors often require some amount of engineering in order to be effective in human cells (Xu et al. 2021; Kannan et al. 2025). Others have taken the alternative approach of minimizing field-standard effectors (e.g. SpCas9 from *S. pyogenes*), reducing overall protein size by rationally deleting structural elements that are non-essential for target binding, specificity and/or editing activity (Shams et al. 2021).

Screening a library of amino acid deletions is an efficient way to identify regions of a protein that may be removed without compromising the protein’s function. However, while there are multiple established mutagenesis techniques which enable the effective construction of substitution libraries, there are relatively few methods for creating deletions in a comprehensive manner (i.e. representing a large subset of all possible deletions, without biases in size or position). Previously, our group has described an approach called MISER (<u>**m**</u>inimization via <u>**i**</u>terative <u>**s**</u>ize-<u>**e**</u>xclusion and <u>**r**</u>ecombination) which can generate deletion libraries encompassing both large and small contiguous in-frame deletions at every position within a gene of interest (Shams et al. 2021). While effective, this method exhibits notable limitations, in particular an inability to capture very small (∼1-2 codon) deletions, as well as relying upon an inefficient mutagenesis technique, plasmid recombineering (Higgins et al. 2017), for a key step in the library assembly process. While recent developments in the field of machine learning have produced AI-assisted protein design tools that may enable the creation of minimized functional versions of proteins via *in silico* structural prediction and innovation (Devkota et al. 2024), the limited efficacy of these approaches at present leaves a need for simpler and more effective experimental methods for constructing deletion libraries in proteins.

Building upon our group’s previous work in developing MISER, here we introduce SABER (<u>**S**</u>pRYgestion <u>**a**</u>nd <u>**b**</u>lunt-<u>**e**</u>nd <u>**r**</u>eligation), a conceptually similar but technically distinct approach which allows for the construction of comprehensive and unbiased deletion libraries with unparalleled speed and ease. Whereas MISER employs plasmid recombineering to introduce restriction sites that can be cut and re-ligated to produce deletions with a two-codon “scar” (Shams et al. 2021), SABER leverages the near-PAMless DNA targeting and nuclease activity of SpRYCas9, an engineered SpCas9 variant (Christie et al. 2023), to generate programmed blunt cuts that are then re-ligated. We applied this technique to the small and structurally divergent Cas9 from *S. aureus* (SaCas9) in order to map the landscape of tolerated deletions using a CRISPRi gene repression screen in bacteria. We found that, as in previous work with SpCas9, multiple domains of the protein can be individually deleted, in whole or in part, while preserving DNA binding activity. We proceeded to make and test SaCas9 constructs with one or more large deletions highlighted in the CRISPRi screen and identified multiple single- and stacked deletion constructs that remained competent for DNA binding, with the smallest functional variants being less than 700 amino acids in length.

## Results

### SABER enables rapid construction of comprehensive and unbiased deletion libraries

To construct the dSaCas9 SABER library, a library of spacer sequences was designed and synthesized as a pool of single-stranded DNA oligomers (ssDNA oligos), each encoding a 20-nucleotide spacer targeting SpRYCas9 cleavage to one of all possible codon-codon junctions in the dSaCas9 gene (Figure 1A). As SpRYCas9 exhibits variable *in vitro* cleavage activity at different PAMs (Walton et al. 2020; Christie et al. 2023), the spacer library was designed such that each codon-codon junction would be targeted from both DNA strands, with the rationale that poor SpRYCas9 cleavage on one strand due to a suboptimal PAM would in most cases be compensated for by a higher rate of cleavage on the other strand due to a better PAM. After obtaining the ssDNA oligo spacer library, the oligos were amplified via PCR and cloned into an sgRNA expression vector to produce a library of sgRNA template plasmids.

**Figure 1:**
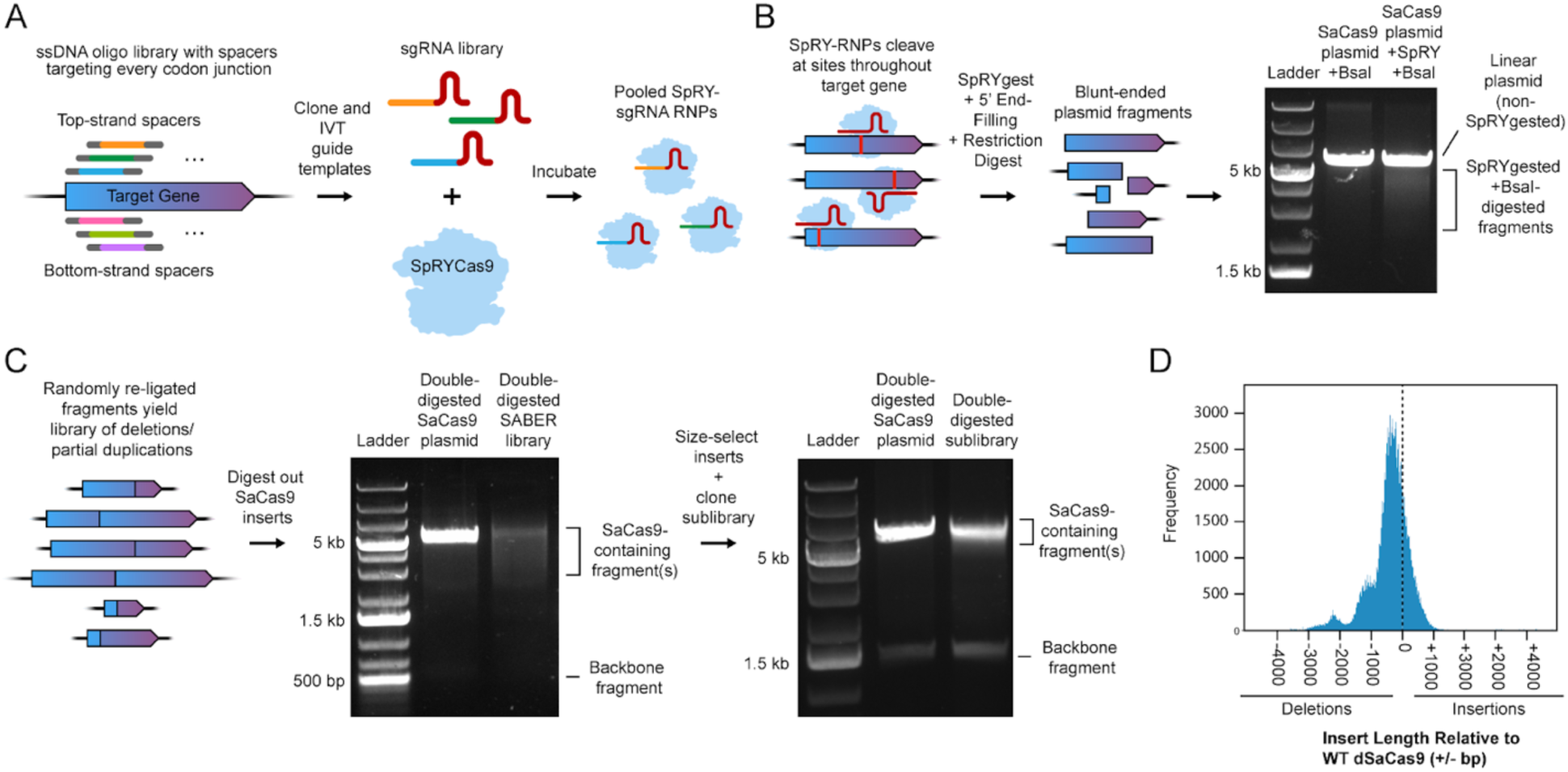
SABER library construction. **A** A library of ssDNA oligos is designed encoding spacers targeting Cas9 cleavage to every inter-codon junction in a gene of interest. Oligos are designed with spacers targeting both top and bottom strands of the target gene. The oligo library is amplified via PCR and cloned into sgRNA expression vectors which are then transcribed to yield a library of sgRNAs. The sgRNAs are incubated with SpRYCas9 to yield SpRY-sgRNA RNPs. **B** SpRYCas9 RNPs digest target gene (in this case dSaCas9) at all inter-codon junctions. Plasmid fragments are treated with T4 DNA polymerase to fill in 5’ overhangs left by SpRYCas9. Fragments are further digested with a restriction enzyme to allow separation of SpRYgested and non-SpRYgested plasmid fragments on an agarose gel. Double-digested fragments are size-selected and extracted. **C** Ligation of extracted fragments yields plasmid library containing deletions and partial duplications within dSaCas9. Plasmids are double-digested to excise dSaCas9 inserts and allow for size-selection of variants likely to be functional (i.e. within ∼500bp of WT dSaCas9). Inserts are recloned into a barcoded expression vector to allow barcode-variant mapping via long-read sequencing. **D** Histogram of SaCas9 variant insert lengths in final size-selected and barcoded dSaCas9 SABER library. Most variants are 0-1000 bp smaller than WT and therefore relatively likely to contain functional deletions (WT dSaCas9 omitted from histogram).

In order to generate the plasmid DNA fragments that would later be ligated to produce the SABER library, the sgRNA library was incubated with purified SpRYCas9 to yield a pool of active SpRY-RNPs, which were then used for *in vitro* digestion of a plasmid encoding the dSaCas9 (i.e. enzymatically dead SaCas9) gene (Figure 1B). By using a sub-stoichiometric ratio (∼1:2) of RNP:plasmid DNA, a level of plasmid cleavage was reached wherein a majority of plasmids were cut by SpRYCas9 either once or not at all, as indicated by subsequent digestion at a restriction site in the plasmid backbone yielding a substantial linear (i.e. having been cut only by the restriction enzyme, and not previously by SpRYCas9) plasmid band on an agarose gel (Figure 1B). This ratio was chosen to minimize bias in deletion size and position, as excessive cutting would result in a high proportion of greatly truncated gene fragments.

Following SpRYCas9 cleavage, the SpRYgest products were treated with T4 DNA Polymerase to repair and blunt 5’ overhangs left by SpRYCas9 (Stephenson et al. 2018), digested with a restriction enzyme cutting in the plasmid backbone, and run on an agarose gel to enable isolation of SpRY-digested plasmid fragments from linear non-SpRY-digested plasmid (Figure 1B). The isolated fragments were then incubated in a ligation reaction to produce a library of plasmids encoding dSaCas9 deletions (Figure 1C). This library was subsequently double-digested to excise the dSaCas9 inserts, which were then size-selected on an agarose gel to isolate variants close in size (within ∼500bp) to WT dSaCas9, thereby enriching for variants likely to contain moderately-sized functional deletions. The isolated inserts were then cloned into barcoded bacterial expression vectors to produce a final dSaCas9 SABER library.

Analysis of long-read sequencing data for the final size-selected SABER library identified 386,930 unique variants, of which approximately 82,270 were in-frame deletions within dSaCas9 (Supplemental Figure 1). This subset accounts for approximately 15% of all possible single contiguous in-frame deletions within dSaCas9 (∼554,404 total possible deletions); for comparison, the MISER library previously used to analyze the deletion landscape of SpCas9 contained 27.5% of all possible SpCas9 deletions. However, given that a majority of deletions were found to be within the desired size range and therefore more likely to be functional (Figure 1D), the complexity of the SABER library was deemed to be more than sufficient for the purposes of constructing an informative deletion map.

### SaCas9 tolerates multiple whole-domain deletions while retaining DNA binding activity

The size-selected dSaCas9 SABER library was assayed for bacterial CRISPRi GFP repression in an RFP/GFP reporter *E. coli* strain (Oakes et al. 2014) using flow cytometry, and the plasmid barcodes of the naive and sorted populations were amplified and sequenced via NGS (Illumina) in order to calculate enrichment values (Figure 2A). In combination with variant-barcode mapping enabled by long-read sequencing of the naive library, this data was used to generate a map of functional dSaCas9 deletions (Figure 2B). Similarly to the previously reported deletion landscape of SpCas9, large deletions (∼50-200 amino acids) were observed to be bounded by domain termini, while smaller deletions (∼1-10 amino acids) were found to be tolerated across most regions of the protein sequence. Much like the SpCas9 deletion map, the deletion map for dSaCas9 highlighted multiple distinct regions where large deletions were tolerated, with the four largest regions encompassing or falling within the REC1, REC3, HNH and RuvC-III domains. Two smaller but well-defined regions were also identified within the WED and TOPO domains.

**Figure 2:**
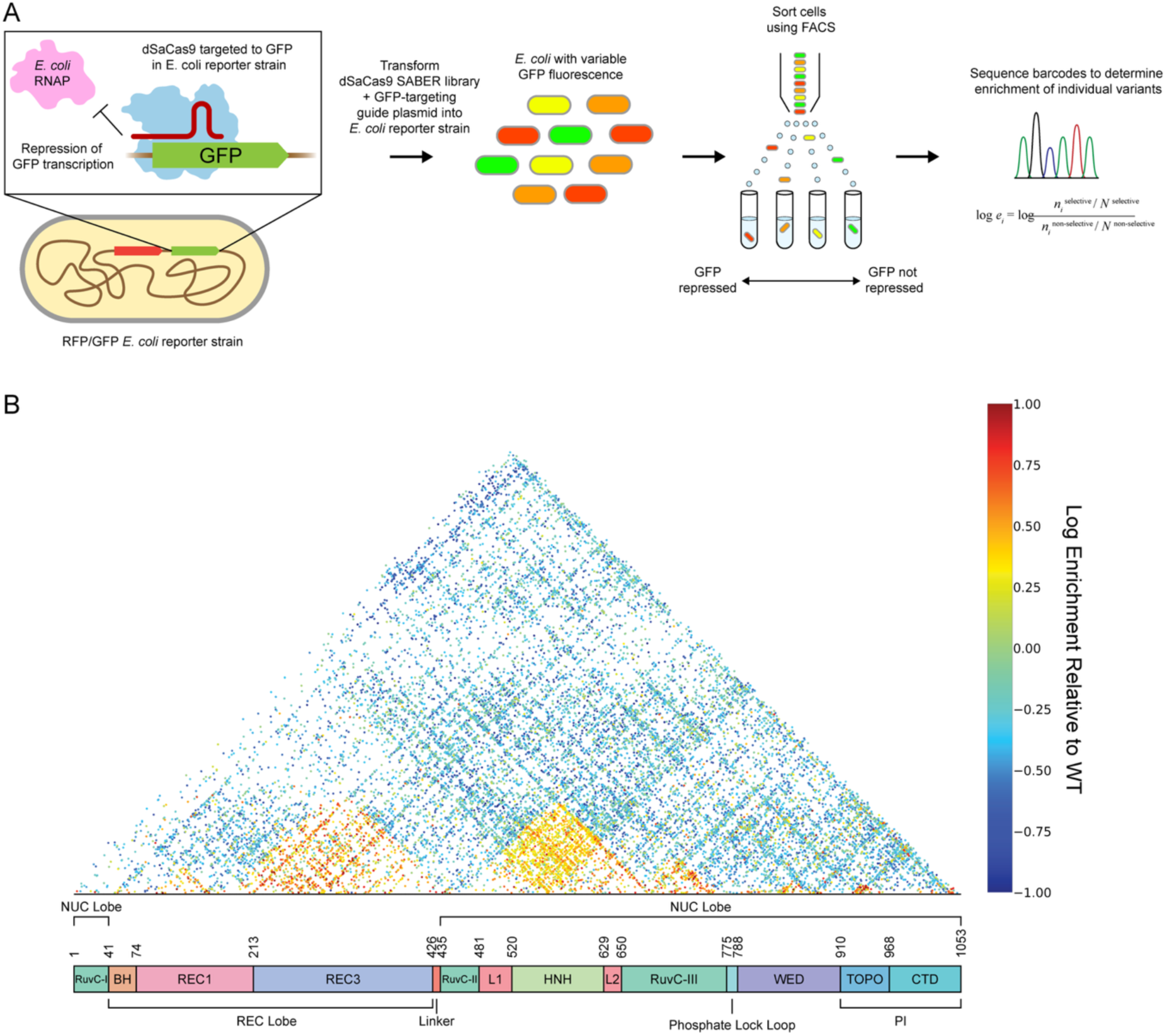
dSaCas9 tolerates large deletions within multiple domains. **A** Schematic of bacterial CRISPRi screen. The dSaCas9 SABER plasmid library is co-transformed into a reporter *E. coli* strain expressing RFP and GFP along with a plasmid expressing an sgRNA targeted to GFP. Expression of functional dSaCas9 variants results in targeted RNP binding to GFP resulting in transcriptional repression. Varying degrees of target-binding result in variable GFP fluorescence enabling sorting of cells containing individual dSaCas9 variants via FACS. Cells are sorted into subpopulations on the basis of GFP/RFP fluorescence ratio. NGS of plasmid barcodes in naive and sorted library populations enables calculation of enrichment values for individual variants. **B** Enrichment map of deletions in dSaCas9. Each colored point sits at the intersection of two diagonal lines which demarcate the N- and C-terminal endpoints of a corresponding deletion where they intersect with the horizontal axis. The color of each point indicates the average enrichment of the associated variant across two replicate transformed and sorted populations taken from a single dSaCas9 SABER library. As can be seen by comparing the distribution of functional deletions to the domain architecture of dSaCas9, most functional deletions occur between the termini of a given domain, with REC1, REC3, HNH, and RuvC-III showing the highest deletion tolerance. The majority of deletions in the library are non-functional, especially those above ∼200 amino acids in size.

Informed by our dSaCas9 deletion map, we proceeded to rationally construct a set of dSaCas9 constructs with large deletions spanning the four major deletion regions highlighted by the enrichment data (Figure 3B). These comprised both constructs with individual deletions (“single-deletion constructs”) as well as constructs with all possible combinations of two or more deletions (“stacked deletion constructs”), for a total of fifteen unique variants. These were assayed individually using a plate-reader fluorescence assay over a 24-hour timecourse, in order to ensure adequate fluorophore maturation and maximal GFP repression at time of final measurement. We found that the single- and stacked deletion constructs exhibited a range of repression activities, with the single-deletion constructs exhibiting repression comparable to WT dSaCas9 and stacked deletion constructs exhibiting several-fold weaker repression than WT dSaCas9.

**Figure 3:**
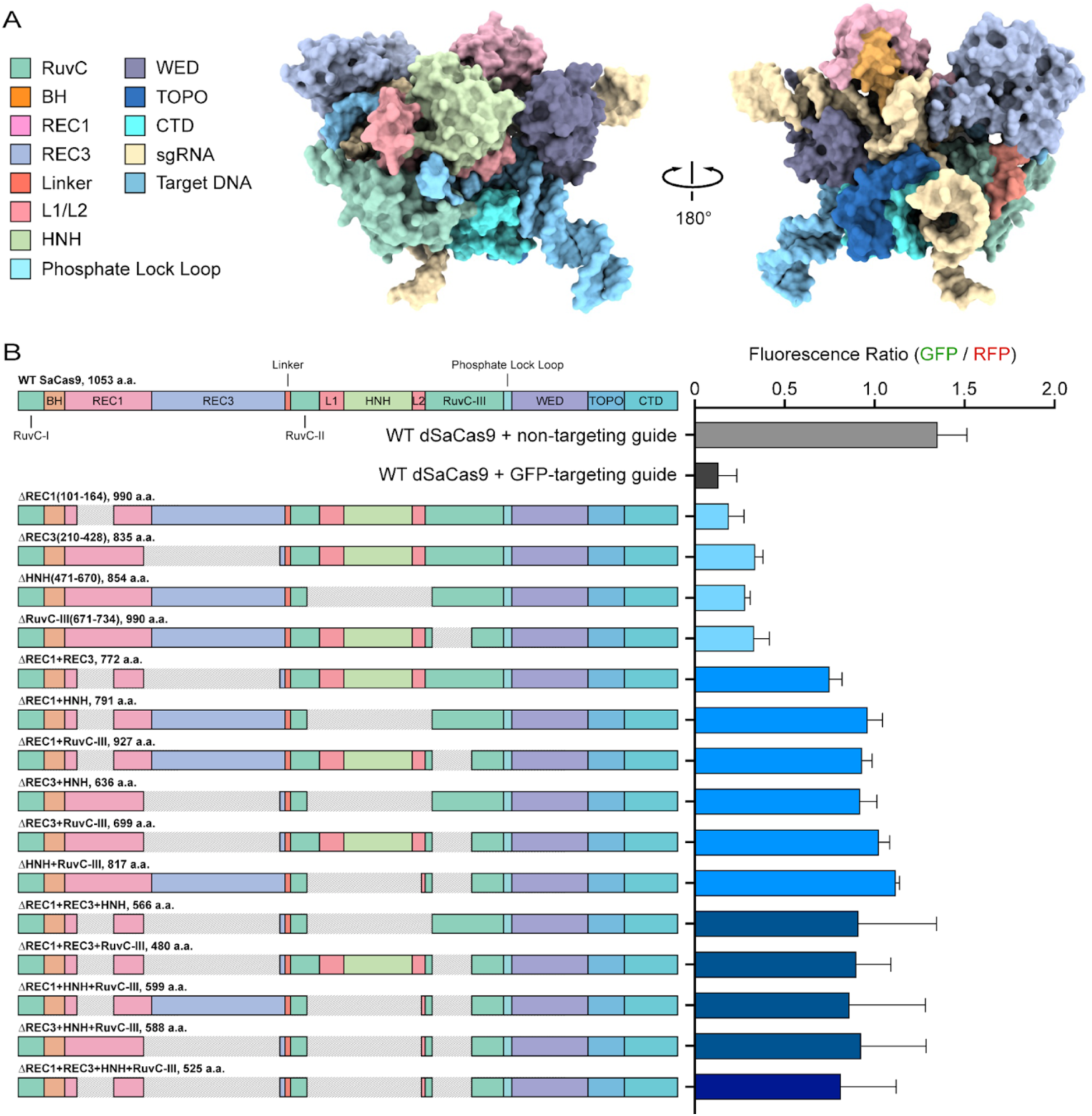
Minimized dSaCas9 constructs show transcriptional repression activity in bacteria. **A** Crystal structure of SaCas9 with annotated domains. **B** dSaCas9 deletion constructs were designed containing rational deletions based on deletion map. Individual testing of dSaCas9 deletion constructs in *E. coli* reporter strain indicates that single-deletion constructs (ΔREC1, ΔREC3, ΔHNH, ΔRuvC-III) exhibit comparable CRISPRi activity to WT dSaCas9. Constructs with multiple deletions show reduced CRISPRi activity.

## Discussion

While multiple well-established DNA engineering techniques exist which enable the random or programmable introduction of substitutions into a gene of interest, there are relatively few options available for making topological changes to a protein’s polypeptide backbone in a similarly high-throughput fashion. Our group’s previous work in developing MISER attempted to address this by introducing a method for programmably constructing unbiased and comprehensive protein deletion libraries of unprecedented depth and complexity, which could be generalized to any protein-coding gene. While highly effective in illuminating the deletion landscape of SpCas9, MISER is unsuitable for investigating a protein’s tolerance for very small (e.g. 1-2 amino acid) deletions - a subset which can potentially have significant effects on protein function, either negative or positive (Patzoldt et al. 2006; Arpino et al. 2014) - as well as relatively time-consuming during the library cloning stage due to the inherent inefficiency of plasmid recombineering (Shams et al. 2021). By comparison, SABER is both simple and fast, with the experimental library construction steps requiring as few as three days and requiring no specialized reagents aside from SpRYCas9 (which is commercially available from NEB) and the ssDNA oligos used to build the SpRY-sgRNA library. While the library constructed for this study contains a relatively high proportion of out-of-frame deletions, we anticipate that this issue can likely be resolved by making minor adjustments to certain steps of the protocol (e.g. 5’ end-filling post-SpRYgestion). As demonstrated above, however, even a library encompassing only ∼15% of all possible in-frame deletions, if unbiased in size and position, can highlight deletion hotspots with fairly high resolution and provide useful insights into the structure-function relationships of a protein.

We chose to investigate the deletion tolerance of SaCas9 in this study principally because of its structural differences from SpCas9, its established utility for *in vivo* AAV delivery and gene editing in human cells, and the reported prevalence of pre-existing immunity against SaCas9 in human subjects (Charlesworth et al. 2019). While our group’s previous work with SpCas9 suggested that multiple domains could likely be individually removed in SaCas9 without complete loss of DNA binding activity, we were initially unsure whether the reduced size and structural divergence of SaCas9 would render it less amenable to large deletions. Our results indicate that despite these differences, SaCas9 tolerates multiple whole- and partial-domain deletions to a similar degree as previously observed with SpCas9.

While SaCas9’s deletion tolerance was found to be broadly similar to SpCas9, we observed unexpected differences in the tolerance for deletions within the REC1 domain. In SpCas9, REC1 is interrupted by an inserted REC2 domain, and while REC2 may be deleted in its entirety without abolishing binding, the two REC1 segments appear to be less tolerant of large deletions (Shams et al. 2021). In contrast, a significant portion of SaCas9’s REC1 domain may be removed with minimal impact on DNA binding (Figure 2B). REC1 exhibits a similar structure in both SpCas9 and SaCas9 (Fig 5A), and in both cases forms contacts with the tracrRNA. Notably, in addition to the large REC2 insertion, the SpCas9 REC1 domain exhibits two smaller insertions which extend out from REC1 to form additional contacts with the tracrRNA repeat-antirepeat duplex (Nishimasu et al. 2015). The large REC1 deletion region highlighted in our SaCas9 deletion map covers a stretch of residues near one of these insertions in the homologous region of the SpCas9 REC1 domain. Given that the large REC1 deletion region highlighted in our SaCas9 deletion map covers a stretch of residues near one of these insertions in the homologous region of the SpCas9 REC1 domain, it may be that this region of REC1 is particularly amenable to indel mutations (ancestrally in the case of SpCas9, as the SpCas9 REC1 domain is currently not especially tolerant of deletions).

**Figure 4:**
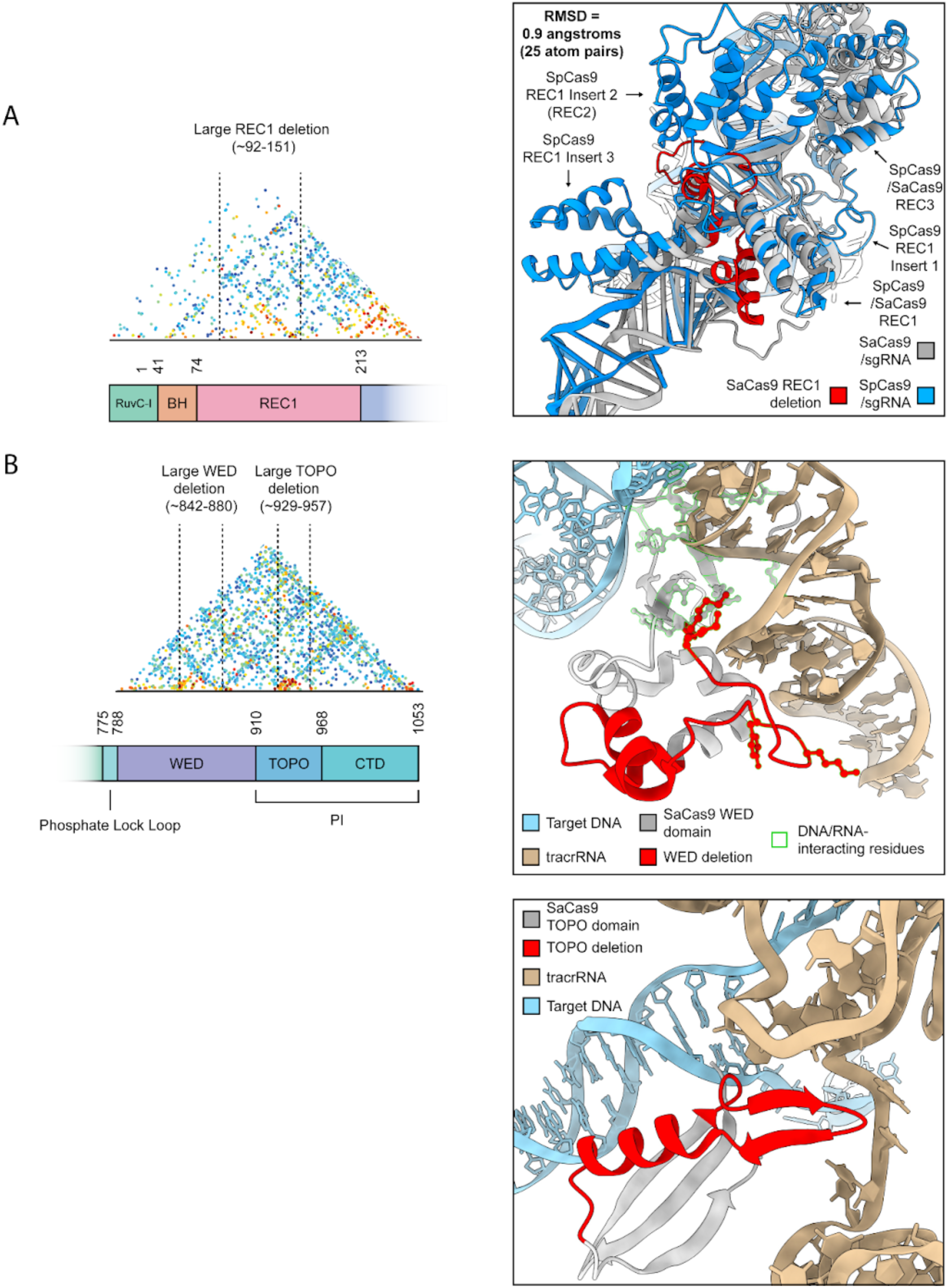
dSaCas9 tolerates small deletions within the WED and TOPO domains. **A** dSaCas9 REC1 tolerates deletions of residues ∼92-151. This deletion spans a segment of REC1 close to the site of an insertion within the homologous REC1 domain in SpCas9. **B** dSaCas9 WED tolerates deletion of residues ∼842-880 while TOPO tolerates deletion of residues ∼929-927. The WED deletion encompasses several tracrRNA-interacting residues while the TOPO deletion removes a motif that appears positioned to interact with tracrRNA.

Also of note are the smaller but well-defined deletion regions within the WED and TOPO domains. The former is a highly-divergent domain recently implicated as a determinant of DNA-unwinding and downstream editing activity in several Cas9 orthologs (Eggers et al. 2024), while the latter forms part of the PAM-Interacting (PI) lobe (Nishimasu et al. 2015). Aside from four residues within the WED deletion region, these deletion regions appear to largely avoid DNA- and RNA-interacting residues within their respective domains, which would seem to emphasize the importance of these residues for tracrRNA and/or target DNA binding (Fig. 5B). However, while the TOPO deletion does not contain any previously-annotated RNA-binding residues, it does encompass a β-sheet motif which projects into a single-stranded loop within the adjacent tracrRNA, with one lysine side chain (Lys935) coming with ∼4 angstroms of the RNA phosphodiester backbone, suggesting a possible role in tracrRNA stabilization. While they are evidently tolerated by dSaCas9 with respect to DNA-binding activity, it remains to be determined whether these WED and TOPO deletions have any subtler impacts on tracrRNA binding.

Previous work has determined that the SaCas9 REC lobe contains a significant number of immunogenic epitopes, with at least 19 residues in this domain having been identified as B-cell antibody binding sites. In particular, mutation of R338 has been demonstrated to reduce antigen-antibody reactivity by around 70%, while mutating E271 and P321 may reduce reactivity by around 60% (Shen et al. 2022). All three of these residues fall within the large REC3 deletion region identified in our deletion map, suggesting that ΔREC3, if demonstrated to be competent for DNA-binding in human cells, may be of utility as a small and minimally immunogenic effector for therapeutic gene-editing applications in humans.

As with MISER before it, SABER can be applied to probe the deletion landscape of any protein-coding gene, so long as a functional screen exists that enables the selection of desired variants. Additionally, the scarless nature of the generated deletions renders SABER suitable for making and screening deletions in non-coding RNAs, where single-nucleotide deletions are broadly more consequential for RNA secondary structure, folding and function than single-amino acid deletions in proteins. We anticipate that SABER’s simplicity and ease of use may encourage its adoption by other groups who seek to minimize proteins with size-sensitive biotechnology applications, including but not limited to additional RNA-guided effector modules and other therapeutic AAV protein cargos such as recombinant CFTR (Zhang et al. 2009). Likewise, we expect that SABER may prove useful as a tool for investigating structure-function relationships in proteins, as well as potentially in large and complex functional RNAs (e.g. ribosomal RNAs).

## Materials and Methods

### Molecular Biology

All restriction enzymes were purchased from New England Biolabs (NEB). PCR was performed using Q5 High-Fidelity DNA Polymerase from NEB. Ligation was performed using T4 DNA Ligase from NEB. Agarose gel extraction was performed using the Zymoclean Gel DNA Recovery Kit, and PCR clean-up was performed using the “DNA Clean & Concentrator” kit, both from Zymo Research. Plasmids were isolated using the QIAprep Spin Miniprep Kit from Qiagen. All DNA-modifying procedures were performed according to the manufacturers’ instructions.

### SABER Library Construction: sgRNA Library Design and Synthesis

A custom Python script was used to generate a library of DNA sequences individually consisting of a spacer designed to target SpRYCas9 to an inter-codon junction in SaCas9, flanked by PCR primer binding sites (Supplementary Table 1). In order to ensure that no cut sites would be excluded by virtue of being associated with an incompatible PAM, spacers were designed with complementarity to both top and bottom DNA strands, with each inter-codon junction being targeted for cleavage from both sides. An ssDNA oligomer library encoding these sequences was ordered from Twist Bioscience, amplified via PCR using primers containing Type IIS restriction sites, and cloned into an sgRNA expression vector to generate a library of sgRNA template plasmids. Template plasmids were linearized via restriction digest and used to generate sgRNAs via *in vitro* transcription with T7 RNA polymerase as described in Walton et al. 2020.

### SABER Library Construction: SpRYgests, Library Cloning and Size-Selection

SpRYCas9 was purified as described in Shams et al. 2021, and complexed with purified library sgRNAs to form SpRY-sgRNA RNPs as described in Walton et al. 2020. To generate programmed cuts at inter-codon junctions in the dSaCas9 gene, purified SpRYCas9 was incubated with *in vitro* transcribed library sgRNAs as described in Walton et al. 2020, and added to restriction digest reactions with dSaCas9 plasmids (typically 2-5 ug total DNA, 0.03uM DNA per each 100uL reaction) at an RNP:DNA ratio of ∼1:2 (∼0.37 uM SpRY-RNP, 0.03 uM circular plasmid DNA). SpRYgests were incubated at 37 °C for 4 hours, and DNA cleavage products were subsequently cleaned using Zymo-Spin II columns and added to a 5’ end-filling reaction with T4 DNA Polymerase (12 °C, 15 minutes). End-filling products were cleaned using Zymo-Spin I columns and added to a BsaI-HfV2 digest reaction (37 °C, 1 hour). BsaI digest products were run on an agarose gel (0.8% agarose-TAE), and SpRYgested fragments were isolated from non-SpRYgested linear plasmid via manual excision and gel extraction. Extracted fragments (typically 0.5-1 ug total DNA) were added to T4 ligation reactions (20 uL reactions, 100 ng DNA each) and incubated at 16 °C overnight.

The following day, ligation products were cleaned using Zymo-Spin I columns and transformed into Stable electrocompetent *E. coli* (New England Biolabs) via electroporation. Transformed cells were transferred to a flask containing 100 mL of media (2xyT + 77.4 µM chloramphenicol) and incubated at 30 °C in a shaking incubator until reaching an OD of ∼1.0, at which point the library plasmids were extracted via miniprep. 3 ug of circular library plasmid DNA was added to a double restriction digest with PaqCI and BsaI-HfV2 (37 °C, 2 hours) and subsequently run on an agarose gel to enable size-selection of dSaCas9 inserts via manual excision and gel extraction.

Barcoded destination vectors were prepared using the NEBuilder HiFi DNA assembly kit to perform ssDNA bridging assembly of a digested vector using ssDNA oligos containing 20 nucleotides of random sequence flanked by 50-nucleotide homology arms. Assembly reactions were transformed into Stable electrocompetent *E. coli* via electroporation. Transformed cells were added to a flask containing 50mL of selective media (2xyT + 237 uM carbenicillin) and grown in a shaking incubator at 37 °C until reaching OD = ∼0.6. Plasmids were harvested via miniprep. Transformation efficiency was estimated by diluting by serially diluting and plating a sample of culture (typical efficiency was > 1E7 CFU).

After assembling the barcoded plasmids, the total extracted dSaCas9 insert DNA was added to a Golden Gate Assembly reaction with BsaI-HfV2, T7 DNA Ligase and 75 ng of PaqCI-digested barcoded destination vector (70 cycles: 5 minutes 37 °C → 5 minutes 16 °C; final 1 hour 16 °C step followed by 5 minutes 60 °C heat inactivation), cleaned using Zymo-Spin I columns, and transformed into Stable electrocompetent *E. coli* via electroporation. Transformed cells were serially diluted (1x, 10x, 100x, 1000x dilution), and 2 mL of each dilution was spread using glass beads on two large square bioassay plates (245 × 245 × 25 mm) with selective agar media (LB agar + 237 uM carbenicillin) and incubated overnight at 30 °C.

CFU counts for each plate were estimated the following day using an optical colony counter, and an area of each plate was manually scraped using a cell lifter to transfer roughly 100,000 colonies to flasks containing 100mL of media (LB + 237 uM carbenicillin). Barcoded library plasmids were extracted via miniprep and stored at -20 °C for subsequent long-read sequencing (PacBio) and use in the fluorescence repression screen.

### Fluorescence Repression Assays: Flow Cytometry

The dSaCas9 SABER library variants were co-transformed via electroporation into a GFP/RFP *E. coli* reporter strain along with a plasmid encoding a GFP-targeting sgRNA as previously described. Transcription of the GFP-targeting sgRNA in conjunction with expression of a functional DNA-binding dSaCas9 variant or WT dSaCas9 results in repression of GFP fluorescence. dSaCas9 was expressed via arabinose induction (araBAD promoter, 0.5 mM arabinose in media). Cells were grown in liquid media (LB + 0.5 mM arabinose + 123.8 µM kanamycin + 77.4 µM chloramphenicol + 237 µM carbenicillin) in a shaking 37 °C incubator to an OD of approximately 0.6 and stored at 4 °C overnight in preparation for sorting the following day. Immediately prior to sorting, cells were diluted in 1X PBS and stored on ice until sorting.

Sorting was performed on a BD FACSAria Fusion Flow Cytometer (BD Biosciences). Cells were sorted on the basis of measured GFP fluorescence relative to RFP (Supplemental Figure 2); briefly, gates were drawn to isolate single *E. coli* cells expressing RFP at near-WT levels, which were then sorted into four bins based on observed GFP/RFP fluorescence ratio. Cell populations from two replicate transformations of the same dSaCas9 SABER library were sorted into PBS (∼0.5 - 2 mL) and subsequently plated using glass beads on large square bioassay plates (two plates per sorted population sample, eight plates total per replicate) with selective agar media (LB agar + 237 µM carbenicillin + 77.4 µM chloramphenicol + 123.8 µM kanamycin) and incubated at 30 °Covernight. The next day, plates were manually scraped to transfer all colonies from a given sorted population into flask containing 100mL of selective media (LB + 237 µM carbenicillin + 77.4 µM chloramphenicol + 123.8 µM kanamycin), and plasmids were subsequently extracted via miniprep.

### Fluorescence Repression Assays: Next-Generation Sequencing

The linearized naive SABER plasmid library was sequenced on a PacBio Revio long-read sequencing device (19,986,902 reads). Sequencing analysis was performed using custom Python scripts (scripts available on request).

The barcode-containing regions of the plasmids isolated from the FACS-sorted *E. coli* populations were amplified via PCR and further processed for Illumina sequencing by the Innovative Genomics Institute NGS Core. Sequencing analysis was performed using custom Python scripts.

### Fluorescence Repression Assays: Validating Isolated Single- and Stacked Deletion Constructs

Rationally-designed single- and stacked deletion constructs were designed as described above and cloned using Golden Gate assembly. Deletion construct plasmids were individually transformed into our GFP/RFP *E. coli* reporter strain alongside a GFP guide plasmid as described previously and plated on solid agar media (LB + arabinose + 237 µM carbenicillin + 77.4 µM chloramphenicol + 123.8 µM kanamycin). Individual colonies were picked and used to inoculate cultures (100uL volume, LB + 237 µM carbenicillin + 77.4 µM chloramphenicol + 123.8 µM kanamycin) in a 96-well plate. GFP, RFP and OD600 measurements were taken over the course of 24 hours using a Tecan 96-well plate reader.

## Acknowledgements

We thank Naiya Phillips (University of California, Berkeley, CA) for her assistance with Cas9 protein purification.

## References

Anzalone, Andrew V., Xin D. Gao, Christopher J. Podracky, Andrew T. Nelson, Luke W. Koblan, Aditya Raguram, Jonathan M. Levy, Jaron A. M. Mercer, and David R. Liu. 2021. “Programmable Deletion, Replacement, Integration and Inversion of Large DNA Sequences with Twin Prime Editing.” Nature Biotechnology. 10.1038/s41587-021-01133-w.

Arpino, James A. J., Samuel C. Reddington, Lisa M. Halliwell, Pierre J. Rizkallah, and D. Dafydd Jones. 2014. “Random Single Amino Acid Deletion Sampling Unveils Structural Tolerance and the Benefits of Helical Registry Shift on GFP Folding and Structure.” Structure (London, England: 1993) 22 (6): 889–98.

Burstein, David, Lucas B. Harrington, Steven C. Strutt, Alexander J. Probst, Karthik Anantharaman, Brian C. Thomas, Jennifer A. Doudna, and Jillian F. Banfield. 2017. “New CRISPR-Cas Systems from Uncultivated Microbes.” Nature 542 (7640): 237–41.

Charlesworth, Carsten T., Priyanka S. Deshpande, Daniel P. Dever, Joab Camarena, Viktor T. Lemgart, M. Kyle Cromer, Christopher A. Vakulskas, et al. 2019. “Identification of Preexisting Adaptive Immunity to Cas9 Proteins in Humans.” Nature Medicine 25 (2): 249–54.

Christie, Kathleen A., Jimmy A. Guo, Rachel A. Silverstein, Roman M. Doll, Megumu Mabuchi, Hannah E. Stutzman, Jiecong Lin, et al. 2023. “Precise DNA Cleavage Using CRISPR-SpRYgests.” Nature Biotechnology 41 (3): 409–16.

Devkota, Kapil, Daichi Shonai, Joey Mao, Scott Soderling, and Rohit Singh. 2025. “Miniaturizing, Modifying, and Augmenting Nature’s Proteins with Raygun.” Preprint (Biorxiv). doi:10.1101/2024.08.13.607858

Doudna, Jennifer A. 2020. “The Promise and Challenge of Therapeutic Genome Editing.” Nature. Nature Research. 10.1038/s41586-020-1978-5.

Eggers, Amy R., Kai Chen, Katarzyna M. Soczek, Owen T. Tuck, Erin E. Doherty, Bryant Xu, Marena I. Trinidad, Brittney W. Thornton, Peter H. Yoon, and Jennifer A. Doudna. 2024. “Rapid DNA Unwinding Accelerates Genome Editing by Engineered CRISPR-Cas9.” Cell 187 (13): 3249-3261.e14.

Gregory, T. Ryan. 2004. “Insertion-Deletion Biases and the Evolution of Genome Size.” Gene. Elsevier. 10.1016/j.gene.2003.09.030.

Higgins, Sean A., Sorel V. Y. Ouonkap, and David F. Savage. 2017. “Rapid and Programmable Protein Mutagenesis Using Plasmid Recombineering.” ACS Synthetic Biology 6 (10): 1825–33.

Hwang, Junghyun, Kwang-Hwi Cho, Han Song, Hyojeong Yi, and Heenam Stanley Kim. 2014. “Deletion Mutations Conferring Substrate Spectrum Extension in the Class A β-Lactamase.” Antimicrobial Agents and Chemotherapy 58 (10): 6265–69.

Kannan, Soumya, Han Altae-Tran, Shiyou Zhu, Peiyu Xu, Daniel Strebinger, Rachel Oshiro, Guilhem Faure, et al. 2025. “Evolution-Guided Protein Design of IscB for Persistent Epigenome Editing in Vivo.” Nature Biotechnology, May. 10.1038/s41587-025-02655-3.

Nishimasu, Hiroshi, Le Cong, Winston X. Yan, F. Ann Ran, Bernd Zetsche, Yinqing Li, Arisa Kurabayashi, Ryuichiro Ishitani, Feng Zhang, and Osamu Nureki. 2015. “Crystal Structure of Staphylococcus Aureus Cas9.” Cell 162 (5): 1113–26.

Oakes, Benjamin L., Dana C. Nadler, and David F. Savage. 2014. “Protein Engineering of Cas9 for Enhanced Function.” Methods in Enzymology 546 (C): 491–511.

Patzoldt, William L., Aaron G. Hager, Joel S. McCormick, and Patrick J. Tranel. 2006. “A Codon Deletion Confers Resistance to Herbicides Inhibiting Protoporphyrinogen Oxidase.” Proceedings of the National Academy of Sciences of the United States of America 103 (33): 12329–34.

Savino, Simone, Tom Desmet, and Jorick Franceus. 2022. “Insertions and Deletions in Protein Evolution and Engineering.” Biotechnology Advances. Elsevier Inc. 10.1016/j.biotechadv.2022.108010.

Shams, Arik, Sean A. Higgins, Christof Fellmann, Thomas G. Laughlin, Benjamin L. Oakes, Rachel Lew, Shin Kim, et al. 2021. “Comprehensive Deletion Landscape of CRISPR-Cas9 Identifies Minimal RNA-Guided DNA-Binding Modules.” Nature Communications 12 (1): 5664.

Shen, Xiaoting, Qinru Lin, Zhiming Liang, Jing Wang, Xinyi Yang, Yue Liang, Huitong Liang, et al. 2022. “Reduction of Pre-Existing Adaptive Immune Responses against SaCas9 in Humans Using Epitope Mapping and Identification.” The CRISPR Journal 5 (3): 445–56.

Stephenson, Anthony A., Austin T. Raper, and Zucai Suo. 2018. “Bidirectional Degradation of DNA Cleavage Products Catalyzed by CRISPR/Cas9.” Journal of the American Chemical Society 140 (10): 3743–50.

Taylor, Martin S., Chris P. Ponting, and Richard R. Copley. 2004. “Occurrence and Consequences of Coding Sequence Insertions and Deletions in Mammalian Genomes.” Genome Research 14 (4): 555–66.

Walton, Russell T., Kathleen A. Christie., Madelynn N. Whittaker, and Benjamin P. Kleinstiver. 2020. “Unconstrained genome targeting with near-PAMless engineered CRISPR-Cas9 variants.” Science 368 (6488), 290–296. doi: 10.1126/science.aba8853

Weiner, January, Francois Beaussart, and Erich Bornberg-Bauer. 2006. “Domain Deletions and Substitutions in the Modular Protein Evolution.” The FEBS Journal 273 (9): 2037–47.

Xu, Xiaoshu, Augustine Chemparathy, Leiping Zeng, Hannah R. Kempton, Stephen Shang, Muneaki Nakamura, and Lei S. Qi. 2021. “Engineered Miniature CRISPR-Cas System for Mammalian Genome Regulation and Editing.” Molecular Cell 81 (20): 4333-4345.e4.

Yang, Gloria, Dave W. Anderson, Florian Baier, Elias Dohmen, Nansook Hong, Paul D. Carr, Shina Caroline Lynn Kamerlin, Colin J. Jackson, Erich Bornberg-Bauer, and Nobuhiko Tokuriki. 2019. “Higher-Order Epistasis Shapes the Fitness Landscape of a Xenobiotic-Degrading Enzyme.” Nature Chemical Biology 15 (11): 1120–28.

Yi, Hyojeong, Karan Kim, Kwang-Hwi Cho, Oksung Jung, and Heenam Stanley Kim. 2012. “Substrate Spectrum Extension of PenA in Burkholderia Thailandensis with a Single Amino Acid Deletion, Glu168del.” Antimicrobial Agents and Chemotherapy 56 (7): 4005–8.

Zhang, Liqun, Brian Button, Sherif E. Gabriel, Susan Burkett, Yu Yan, Mario H. Skiadopoulos, Yan Li Dang, et al. 2009. “CFTR Delivery to 25% of Surface Epithelial Cells Restores Normal Rates of Mucus Transport to Human Cystic Fibrosis Airway Epithelium.” PLoS Biology 7 (7): e1000155.

