## Supplementary Information for "Exploring the deletion landscape of *S. aureus* Cas9 with SABER"

**Andrew J. Plebanek et al. 2025,**

**Supplementary Information**

| Oligo | Purpose | Sequence |
|---|---|---|
| osgRNA_TOP(*n*) (generic sequence) | sgRNA library spacer oligo (top strand spacers) | GGGTAGTCATGTGACGAAGTACCGATGCCACACGGTCTCATAGG(placeholder: 20-nt spacer derived from dSaCas9)GTTTAGAGACCCACACCACTGACGCTGATACCGAAGCGGGCTTATA |
| osgRNA_BOT(*n*) (generic sequence) | sgRNA library spacer oligo (bottom strand spacers) | CAGCGAAAAGGTGACGAAGTACCGATGCCACACGGTCTCATAGG(placeholder: 20-nt spacer derived from dSaCas9)GTTTAGAGACCCACACCACTGACGCTGATACCGAAGCCCAGTATTC |
| oAJP472 | Forward PCR primer for amplifying sgRNA spacer library oligos | GTGACGAAGTACCGATGC |
| oAJP473 | Reverse PCR primer for amplifying sgRNA spacer library oligos | CTTCGGTATCAGCGTCAG |
| oAJP774 | Barcoded oligo for ssDNA bridging assembly barcoding reaction | CAAAACGATCTCAAGAAGATCATCTTATTAATCAGATAAAATATTTCTAGNNNNNNNNNNNNNNNNNNNNNCTAGATTTCAGTGCAATTTATCTCTTCAAATGTAGCACCTGAAGTCAGCC |
| oBB1_FW | Forward PCR primer for generating N-terminal dSaCas9 fragments to be used when cloning dSaCas9 | CACACCAGGTCTCAgcgccgcagtaacaattg |

| | | |
|---|---|---|
| | deletion constructs (paired with DR(*n*)_RV primers) | |
| oBB1_RV | Reverse PCR primer for generating C-terminal dSaCas9 fragments to be used when cloning dSaCas9 deletion constructs (paired with DR(*n*)_RV primers) | CACACCAGGTCTCAgcgcatggaagcgattaa |
| oDR1_FW2 | Forward PCR primer for cloning REC1 deletion (paired with BB1_RV) | CACACCAGGTCTCACGGCcggggcagcatcaacaga |
| oDR1_RV1 | Reverse PCR primer for cloning REC1 deletion (paired with BB1_FW) | CACACCAGGTCTCAGCCGCCctcgctcagcttctggct |
| oDR2_FW3 | Forward PCR primer for cloning REC3 deletion (paired with BB1_RV) | CACACCAGGTCTCACGGCgtggacctgtcccagcag |
| oDR2_RV3 | Reverse PCR primer for cloning REC3 deletion (paired with BB1_FW) | CACACCAGGTCTCAGCCGCCggtccgccgggtttccag |
| oDR3_FW2 | Forward PCR primer for cloning HNH deletion (paired with BB1_RV) | CACACCAGGTCTCACGGCgtgaaagtgaagtccatc |
| oDR3_RV2 | Reverse PCR primer for cloning HNH deletion (paired with BB1_FW) | CACACCAGGTCTCAGCCGCCcaggccgtacttcttgat |
| oDR4_FW2 | Forward PCR primer for cloning RuvC-III deletion (paired with BB1_RV) | CACACCAGGTCTCACGGCgaggaaaagcaggccgag |
| oDR4_RV2 | Reverse PCR primer for cloning RuvC-III deletion (paired with BB1_FW) | CACACCAGGTCTCAGCCGCCcaggttgttcactctgaa |
| oCsiI_Primer | Primer with CsiI site to enable cloning of deletion constructs with stacked | CACACCAaccaggttGCCGCCcaggccgtacttcttgat |

| | |
|---|---|
| | HNH+RuvC-III deletions (paired with BB1_FW) |

**Supplemental Table 1: ssDNA oligomers used in this study.**

| Plasmid | Purpose |
|---|---|
| pCIT101 | Bacterial dSaCas9 part vector |
| pCIT116 | Bacterial destination vector: dSaCas9 insert |
| pAJP143 | Bacterial protein expression vector: WT dSaCas9 (control) |
| pAJP082 | Bacterial destination vector: SpRYCas9 sgRNA spacer acceptor |
| pCIT114 | Bacterial sgRNA expression vector: GFP-targeting SaCas9 sgRNA |
| pAJP141 | Bacterial sgRNA expression vector: Hemoglobin-targeting SaCas9 sgRNA (control) |
| pAJP145 | Bacterial protein expression vector: ΔREC1 dSaCas9 construct |
| pAJP146 | Bacterial protein expression vector: ΔREC3 dSaCas9 construct |
| pAJP147 | Bacterial protein expression vector: ΔHNH dSaCas9 construct |
| pAJP148 | Bacterial protein expression vector: ΔRuvC-III dSaCas9 construct |
| pAJP149 | Bacterial protein expression vector: ΔREC1+REC3 dSaCas9 construct |
| pAJP150 | Bacterial protein expression vector: ΔREC1+HNH dSaCas9 construct |
| pAJP151 | Bacterial protein expression vector: ΔREC1+RuvC-III dSaCas9 construct |
| pAJP152 | Bacterial protein expression vector: ΔREC3+HNH dSaCas9 construct |
| pAJP153 | Bacterial protein expression vector: ΔREC3+RuvC-III dSaCas9 construct |
| pAJP154 | Bacterial protein expression vector: ΔHNH+RuvC-III dSaCas9 construct |
| pAJP155 | Bacterial protein expression vector: ΔREC1+REC3+HNH dSaCas9 construct |
| pAJP156 | Bacterial protein expression vector: ΔREC3+HNH+RuvC-III dSaCas9 construct |

| | |
|---|---|
| pAJP157 | Bacterial protein expression vector: ΔREC1+HNH+RuvC-III dSaCas9 construct |
| pAJP158 | Bacterial protein expression vector: ΔREC1+REC3+RuvC-III dSaCas9 construct |
| pAJP159 | Bacterial protein expression vector: ΔREC1+REC3+HNH+RuvC-III dSaCas9 construct |

**Supplemental Table 2: Plasmids used in this study.**

| | |
|---|---|
| **Polymerase read bases** | 1,319,237,465,200 |
| **Polymerase reads** | 19,986,902 |
| **Polymerase read length (mean)** | 66.01 kb |
| **Polymerase read length (N50)** | 126.25 kb |
| **Polymerase read length longest subread length (mean)** | 8.42 kb |
| **Polymerase read length longest subread length (N50)** | 8.25 kb |
| **Unique molecular yield** | 151,466,967,040 |
| **Local base rate** | 2.23 |

**Supplemental Table 3: PacBio long-read sequencing statistics for the naive size-selected and barcoded dSaCas9 SABER library.**

| **Sample** | **Read count** | **Read quality (mean errQ of forward and reverse read set)** |
|---|---|---|
| Naive library (pre-transformation) | 6843652 | 39.1 |
| Library transformation replicate A (starting population/pre-selection) | 7561000 | 39.1 |
| Library transformation replicate B (starting population/pre-selection) | 6529667 | 39.15 |
| Sorted population: highest GFP repression (replicate A) | 7064298 | 39.15 |
| Sorted population: highest GFP repression (replicate B) | 9881943 | 39.1 |

| | | |
|---|---|---|
| Sorted population: second-highest GFP repression (replicate A) | 8398569 | 39.15 |
| Sorted population: second-highest GFP repression (replicate B) | 11253616 | 39.1 |
| Sorted population: third-highest GFP repression (replicate A) | 6892878 | 39.15 |
| Sorted population: third-highest GFP repression (replicate B) | 8166489 | 39.2 |
| Sorted population: no GFP repression (replicate A) | 6775474 | 39.1 |
| Sorted population: no GFP repression (replicate B) | 8954964 | 39.15 |

**Supplemental Table 4: Illumina sequencing statistics for naive and sorted dSaCas9 SABER libraries.**

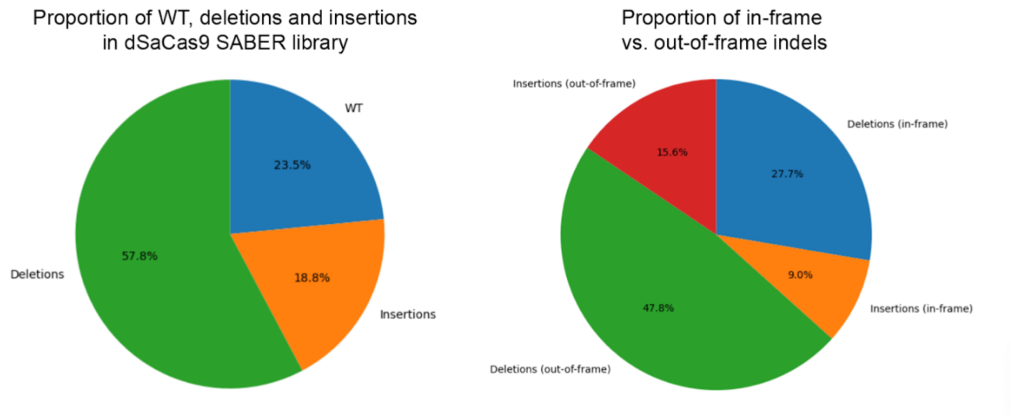

**Supplemental Figure 1: Variant-calling statistics for PacBio-sequenced naive size-selected and barcoded dSaCas9 SABER library.** Percentages indicate proportion of mapped unique barcode-variant pairs falling into each category.

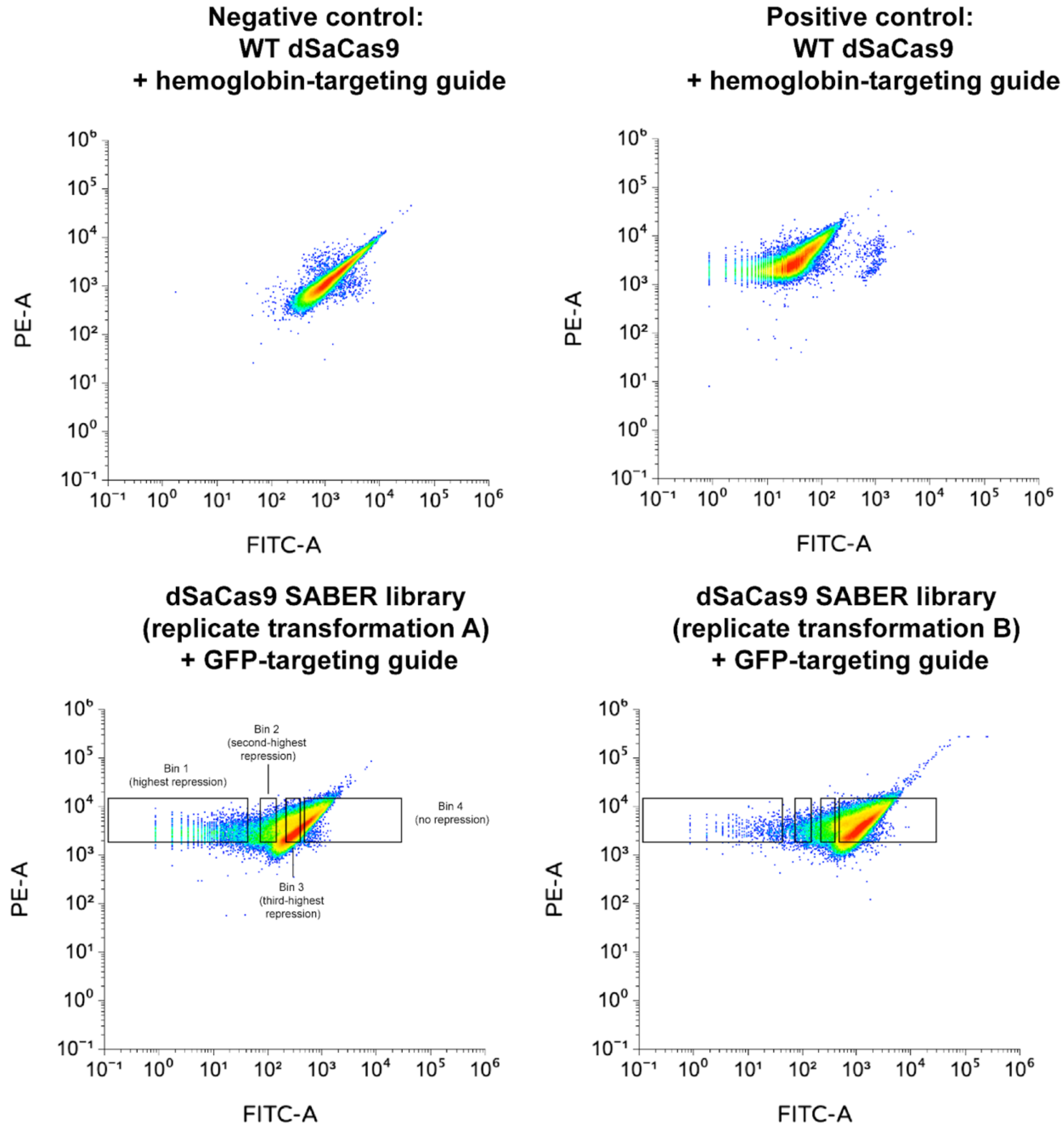

**Supplemental Figure 2: FACS traces for controls and library samples analyzed in dSaCas9 CRISPRi FACS experiment.** PE (phycoerythrin, excitation peak = 565 nm, emission peak = 578 nm) is here used as a calibration proxy for mRFP1 (monomeric red fluorescent protein from *Discosoma* sp., excitation peak = 590 nm, emission peak = 612 nm), while FITC (fluorescein isothiocyanate, excitation peak = 498 nm, emission peak = 517 nm) is used as a calibration proxy for sfGFP (Superfolder green fluorescent protein engineered from *Aequorea victoria* GFP, excitation peak = 488 nm, emission peak = 510 nm). Cells were first selected on the basis of forward-scatter (FSC) versus side-scatter (SSC) ratio as well as peak forward scatter intensity (FSC-H) versus total forward scatter (FSC-A) to isolate single cells as opposed to clusters. A subset of this filtered subpopulation was then selected on the basis of having RFP fluorescence values around or above the mean, and then sorted on the basis of GFP fluorescence using four gates (annotated gates in bottom left panel).

| | | |
|---|---|---|
| Experiment : 11_21_24_MISERsort<br>Specimen : Specimen_001<br>Tube : SpikeLibA<br>Sort Layout : Sort Layout_003<br>Application : FACSDiva Version 8.0.3 | Sort Report | Report Date : 2024.11.21 at 22:37:45<br>Device : 4 Tube<br>User ID : Administrator<br>Cytometer : FACSAriaIII (R656700230002) |

**Sort Settings**

| | | | |
|---|---|---|---|
| Sort Setup | 70 micron | Precision | 4-Way Purity |
| Frequency | 87.0 | Yield Mask | 0 |
| Amplitude | 14.6 | Purity Mask | 32 |
| Phase | 0.00 | Phase Mask | 0 |
| Drop Delay | 44.37 | Single Cell | Off |
| Attenuation | Off | Plates Voltage | 5,000 |
| Sweet Spot | On | Voltage Centering | -232 |
| First Drop | 204 | Sheath Pressure | 70.00 |
| Target Gap | 7 | | |

**Side Stream Voltage (%)**

| Far Left | Left | Right | Far Right |
|---|---|---|---|
| 85.00 | 26.00 | 18.00 | 75.00 |

**Neighboring Drop Charge (%)**

| 2nd | 3rd | 4th |
|---|---|---|
| 14.00 | 7.00 | 0.00 |

**Acquisition Counters**

| | |
|---|---|
| Threshold Count | 172176678 |
| Processed Events Count(evt) | 170447888 |
| Electronic Aborts Count(evt) | 5713983 |
| Sort Elapsed Time(hh:mm:ss) | 01:02:51 |

**Sort Counters**

| | Far Left | Left | Right | Far Right |
|---|---|---|---|---|
| Sort Rate(evt/s) | 530 | 29 | 530 | 74 |
| Conflicts Count(evt) | 2849851 | 152123 | 14870038 | 366422 |
| Conflicts Rate(evt/s) | 755 | 40 | 3943 | 97 |
| Efficiency(%) | 41 | 42 | 11 | 43 |

**Sort Layout**

| Far Left | Left | Right | Far Right |
|---|---|---|---|
| P3_NormRFP : 2000000 / 2000000 | P3_NormRFP : 112195 / 2000000 | P3_NormRFP : 2000000 / 2000000 | P3_NormRFP : 281058 / 2000000 |

40

| | | |
|---|---|---|
| Experiment : 11_21_24_MISERsort<br>Specimen : Specimen_001<br>Tube : SpikedLibB<br>Sort Layout : Sort Layout_003<br>Application : FACSDiva Version 8.0.3 | Sort Report | Report Date : 2024.11.21 at 23:57:51<br>Device : 4 Tube<br>User ID : Administrator<br>Cytometer : FACSAriaIII (R656700230002) |

**Sort Settings**

| | | | |
|---|---|---|---|
| Sort Setup | 70 micron | Precision | 4-Way Purity |
| Frequency | 87.0 | Yield Mask | 0 |
| Amplitude | 14.6 | Purity Mask | 32 |
| Phase | 0.00 | Phase Mask | 0 |
| Drop Delay | 44.37 | Single Cell | Off |
| Attenuation | Off | Plates Voltage | 5,000 |
| Sweet Spot | On | Voltage Centering | -232 |
| First Drop | 204 | Sheath Pressure | 70.00 |
| Target Gap | 7 | | |

**Side Stream Voltage (%)**

| Far Left | Left | Right | Far Right |
|---|---|---|---|
| 85.00 | 26.00 | 18.00 | 75.00 |

**Neighboring Drop Charge (%)**

| 2nd | 3rd | 4th |
|---|---|---|
| 14.00 | 7.00 | 0.00 |

**Acquisition Counters**

| | |
|---|---|
| Threshold Count | 171084823 |
| Processed Events Count(evt) | 166198914 |
| Electronic Aborts Count(evt) | 7027811 |
| Sort Elapsed Time(hh:mm:ss) | 00:59:39 |

**Sort Counters**

| | Far Left | Left | Right | Far Right |
|---|---|---|---|---|
| Sort Rate(evt/s) | 329 | 20 | 558 | 81 |
| Conflicts Count(evt) | 1585417 | 110301 | 9966807 | 434228 |
| Conflicts Rate(evt/s) | 442 | 30 | 2784 | 121 |
| Efficiency(%) | 42 | 40 | 16 | 40 |

**Sort Layout**

| Far Left | Left | Right | Far Right |
|---|---|---|---|
| P3_NormRFP : 1180311 / 2000000 | P3_NormRFP : 74112 / 2000000 | P3_NormRFP : 2000000 / 2000000 | P3_NormRFP : 293462 / 2000000 |

41

**Supplemental Figure 3: Sort reports for dSaCas9 CRISPRi FACS experiment.** "Far Left" = highest GFP repression population, "Left" = second-highest GFP repression population, "Right" = non-GFP-repressed population, "Far Right" = third-highest GFP repression population. Readouts generated using BD FACSDIVA™ software.

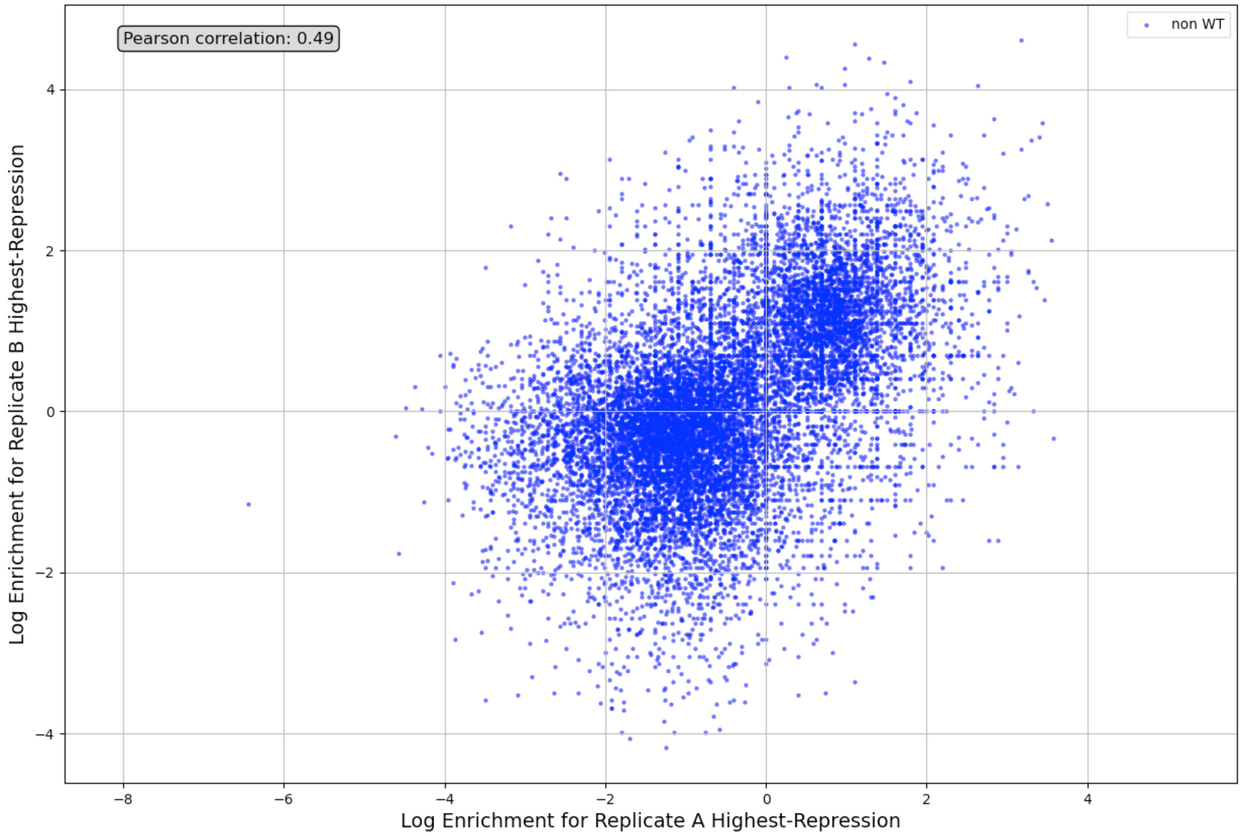

**Supplemental Figure 4: Correlation of log enrichment values between sorted replicate dSaCas9 SABER library transformations for the "Highest-Repression" bin.**

| dSaCas9 variant | **REC1 deletion** | **REC3 deletion** | **HNH deletion** | **RuvC-III deletion** |
|---|---|---|---|---|
| **Δ**REC1 | 101-164 | **-** | **-** | **-** |
| **Δ**REC3 | **-** | 210-428 | **-** | **-** |
| **Δ**HNH | **-** | **-** | 471-670 | **-** |
| **Δ**RuvC-III | **-** | **-** | **-** | 671-734 |
| **Δ**REC1+REC3 | 101-164 | 210-428 | **-** | **-** |
| **Δ**REC1+HNH | 101-164 | **-** | 471-670 | **-** |
| **Δ**REC1+RuvC-III | 101-164 | **-** | **-** | 671-734 |
| **Δ**REC3+HNH | **-** | 210-428 | 471-670 | **-** |
| **Δ**REC3+RuvC-III | **-** | **-** | **-** | 671-734 |
| **Δ**HNH+RuvC-III | **-** | **-** | 471-644 | 671-734 |
| **Δ**REC1+REC3+HNH | 101-164 | 210-428 | 471-670 | **-** |
| **Δ**REC1+REC3+RuvC-III | 101-164 | 210-428 | **-** | 671-734 |
| **Δ**REC1+HNH+RuvC-III | 101-164 | **-** | 471-644 | 671-734 |
| **Δ**REC3+HNH+RuvC-III | **-** | 210-428 | 471-644 | 671-734 |
| **Δ**REC1+REC3 +HNH+RuvC-III | 101-164 | 210-428 | 471-644 | 671-734 |

**Supplemental Table 5: Single- and stacked dSaCas9 deletion constructs.**
